# Absence of the bile acid enzyme CYP8B1 increases brain chenodeoxycholic acid and reduces neuronal excitotoxicity in mice

**DOI:** 10.1101/2022.12.11.520005

**Authors:** Vera F. Monteiro-Cardoso, Xin Yi Yeo, Han-Gyu Bae, David Castano Mayan, Mariam Wehbe, Sejin Lee, Kumar Krishna-K, Seung Hyun Baek, Leon F. Palomera, Sangeetha Shanmugam, Kai Ping Sem, Matthew P. Parsons, Michael R. Hayden, Elisa A. Liehn, Sreedharan Sajikumar, Svend Davanger, Dong-Gyu Jo, Sangyong Jung, Roshni R. Singaraja

**Affiliations:** Yong Loo Lin School of Medicine, National University of Singapore, 117597, Singapore; Translational Laboratories in Genetic Medicine, Agency for Science, Technology and Research, 138648, Singapore; Institute for Molecular and Cellular Biology, Agency for Science, Technology and Research, 138667, Singapore; Department of Psychological Medicine, Yong Loo Lin School of Medicine, National University of Singapore, 119228, Singapore; Department of Life Sciences, Yeungnam University, Gyeongsan 38541, South Korea; Cardiovascular Research Institute, National University Health System, Singapore; Institute of Basic Medical Sciences, University of Oslo, 0317 Norway; Department of Physiology and Healthy Longevity Translational Research Program, Yong Loo Lin School of Medicine, National University Singapore, 117593, Singapore; School of Pharmacy, Sungkyunkwan University, Suwon 16419, Republic of Korea; Division of BioMedical Sciences, Faculty of Medicine, Memorial University of Newfoundland, St. John’s, Newfoundland, A1B 3V6, Canada; Department of Medical Genetics, Centre for Molecular Medicine and Therapeutics, The University of British Columbia, Vancouver, British Columbia, V5Z 4H4, Canada; Victor Babes National Institute of Pathology, Bucharest 050096, Romania; Institute for Molecular Medicine, University of South Denmark, Odense, Denmark

**Keywords:** Bile acid, chenodeoxycholic acid, CYP8B1, excitotoxicity, Ischemic stroke, NMDA receptor, Huntington Disease, excitatory postsynaptic current, glutamate

## Abstract

**Background:** Bile acids (BAs), which act in the liver-brain axis, are liver-derived signaling molecules found in the brain. However, how they modulate neurological function remains largely unknown.

**Methods:** To assess the role of BAs in the brain, we generated mice with absent 12α-hydroxylase (*Cyp8b1*), a BA synthesis enzyme, and determined if brain BA levels were altered in these mice, and if and how this may modulate neuronal function.

**Results:** The absence of CYP8B1 increased brain levels of the primary BA chenodeoxycholic acid (CDCA), and decreased ischemic stroke infarct area. Furthermore, CDCA administration reduced ischemic stroke lesion area in *wild-type* mice. Excitotoxicity due to elevated extra-cellular glutamate contributes to neuronal death in ischemic stroke. Neurons from *Cyp8b1^-/-^* mice showed reduced susceptibility to glutamate-induced toxicity, and exogenous CDCA reduced glutamate-induced toxicity in neurons from *wild-type* mice. These data suggest that CDCA-mediated decreases in excitotoxic neuronal death contributes to the reduced stroke lesion area in *Cyp8b1^-/-^* mice. Aberrant N-methyl-D-aspartate receptor (NMDAR) over-activation contributes to excitotoxicity. CDCA decreased NMDAR-mediated excitatory post-synaptic currents (EPSCs) in *wild-type* brain slices, by reducing over-activation of the NMDAR subunit GluN2B. In line with this, synaptic NMDAR activity was also decreased in *Cyp8b1^-/-^* brain slices. Expression level and synaptic distribution of GluN2B were unaltered in *Cyp8b1^-/-^* mice, suggesting that CDCA may directly antagonize GluN2B-containing NMDARs.

**Conclusions:** Our data suggests that CDCA acts in the liver-brain axis and decreases the aberrant over-activation of neuronal GluN2B-containing NMDARs, contributing to neuroprotection.

## Introduction

The role of the liver in the liver-brain axis has garnered recent attention. One pathway that connects the liver with the brain is the vagus nerve in the autonomic nervous system. Liver derived factors such as bile acids form another pathway linking the brain and liver (1). Bile acids (BAs) are liver-derived amphipathic end products of cholesterol metabolism that aid in the solubilization, digestion and absorption of lipids and lipid-soluble nutrients in the gastrointestinal system (2). The discovery that BAs and their receptors are present in the brain during normal physiological conditions (3-5) raised the possibility that central control of metabolism and neuronal function can be regulated by BAs. Liver-derived BAs can enter the brain through the blood-brain barrier (6), suggesting that BAs may act in the liver-brain axis to modulate neurological function. CYP27A1, a BA synthesis enzyme, is normally present in the brain. Its absence results in brain cholestenol accumulation (7), suggesting that *de novo* BA synthesis may also occur in the brain.

The liver generates two primary BAs, the 12α*-*hydroxylated BA cholic acid (CA), and the non-12α*-*hydroxylated BA chenodeoxycholic acid (CDCA) (**Supplemental Fig 1**) (2). CDCA can be further converted to ursodeoxycholic acid (UDCA) and its taurine-conjugated form TUDCA (2), before entering the bloodstream. CA and CDCA levels are tightly controlled by sterol 12α-hydroxylase (CYP8B1), which synthesizes 12α*-* hydroxylated BAs, including CA (**Supplemental Fig 1**) (8-11). In the absence of CYP8B1, a substantial decrease in CA, and a compensatory increase in CDCA and its non-12α*-*hydroxylated BA derivatives, including TUDCA, are observed (8-11).

Although not much is known about the function of BAs in the brain, administration of TUDCA to mouse models of neurodegenerative disorders reduced their pathologic phenotypes (12). In humans, mutations in the BA synthesis gene *CYP27A1* result in almost absent CDCA synthesis, leading to Cerebrotendinous xanthomatosis (CTX), a rare neurodegenerative disorder (7). Although several BAs, including the intestinal derivatives of CDCA, have been tested as treatments for CTX, CDCA administration is the only effective treatment for the neurological symptoms in patients with CTX (13,14). However, the mechanism by which CDCA reduces CTX symptoms remains unknown. Together, these data suggest that BAs can cross the blood-brain barrier, and may have the potential to directly regulate neuronal function. Thus, further clarification of the functional roles of BAs in neurophysiology are required.

We investigated the role of BAs to act as signalling molecules in the liver-brain axis by assessing the effects of BA dysregulation on neuronal functions in mice with deleted *Cyp8b1*, a predominantly liver expressed BA synthesis enzyme. We find that brain CDCA levels are increased in these mice, and that CDCA protects against neuronal dysfunction by reducing NMDAR-mediated neurotoxicity.

## Methods

Complete information for all materials and methods is provided in the supplemental data.

### Mouse lines and antibiotic treatment

All mouse experiments were approved by the Biomedical Sciences Institute, Singapore, institutional animal care committee, and received care as outlined in the Guide for the Care and Use of Laboratory Animals. Generation of the *Cyp8b1^-/-^* mice is described in the supplemental data. All experiments utilized male mice, housed in 12h light/dark cycles with *ad libitum* access to water and food (1324_modified, Altromin GmbH & Co.). All animals within a group were randomly assigned, and all analyses were performed during the light cycle, by experimenters blind to the genotypes/treatment group. Study design and reporting followed the ARRIVE guidelines. Sample sizes were determined based on previous studies, and no animals were excluded. Antibiotics were administered to *wild-type* mice as described (15). For CDCA administration, 12-week old male *wild-type C57BL/6N* mice were fed standard diet containing 1% CDCA (w/w) for 7 days. Detailed information is provided in the supplemental data.

### Neuronal cell culture and cytotoxicity assay

Mixed cortical cultures were prepared as described (16). After 14 days, cells were pre-incubated with 30µM CDCA (Sigma) or 100µM TUDCA (Sigma) for 3h, followed by 30µM L-glutamate. Cytotoxicity was evaluated in the culture media after 24h, using the Pierce LDH Cytotoxicity Assay Kit (Thermo Scientific).

### Bile acid measurements

BAs were analyzed in plasma and brains from mice by Genome BC Proteomics Center, University of Victoria, Canada, as described (17).

### Transient middle cerebral artery occlusion (tMCAO)

tMCAO was carried out as described (18). 24h post-MCAO, brains were isolated and 2mm coronal sections generated. Lesions were identified by staining with 2% 2,3,5-triphenyltetrazolium (TTC, Sigma) and quantified using ImageJ.

### Preparation of brain slices and whole cell patch

Mice were euthanized by cervical dislocation, brains removed, transferred to oxygenated aCSF, sliced horizontally, and slices placed in a recording chamber containing aCSF with 50µM picrotoxin. NMDAR-and AMPAR-mediated currents of the CA1 pyramidal neurons were evoked by stimulation of Schaffer collateral fibers in the stratum radiatum of CA2. NMDAR-mediated currents were measured at +40mV and amplitude was considered 50ms after stimulation. AMPAR-mediated currents were measured at −70mV and amplitude was measured at peak response.

### Functional inhibition of NMDAR-mediated excitatory postsynaptic currents (EPSCs) by extra-cellular CDCA

Whole-cell patch was conducted in hippocampal CA3-CA1 synaptic circuits in brain slices from *wild-type* mice, with cesium-based internal solution. AMPAR-and NMDAR-mediated EPSCs were measured at −70mV and +40mV of holding membrane potentials in CA1 pyramidal neurons with electric stimulation, before and after perfusion with 0.1µM or 1µM of CDCA in aCSF. To avoid effects from residual binding of CDCA, data were collected from a single cell per brain slice.

### Western immunoblotting

Hippocampal and primary neuron protein lysates were resolved on 7.5% SDS-PAGE gels, transferred onto nitrocellulose membranes, and incubated with antibodies against GluN1, GluN2A, GluN2B, and PARP1 (1:1000, Cell Signalling, USA) or calnexin (1:5000, Sigma-Aldrich), followed by HRP-conjugated secondary antibody (Cell Signalling, 1:5000 for calnexin, 1:2000 for others). Signals were detected using Clarity Western ECL (BioRad). Relative densities were calculated using ImageJ.

### Post-embedding immunogold labeling

Mice were perfused with 4% formaldehyde and 0.1% glutaraldehyde in 0.1M sodium phosphate buffer, and hippocampal CA1 pieces were freeze substituted, embedded in Lowicryl, sectioned, and subjected to immunogold labelling, as described (19). Rabbit anti-NMDAR2B antibody (Proteintech, 1:50), was followed by 12nm colloidal gold-conjugated goat anti-Rabbit IgG (Abcam, 1:20). Transmission electron micrographs of CA1 Schaffer collateral synapses were obtained at 43000x magnification (Tecnai G2 Spirit, FEI Company). Membrane gold particle densities were quantified with ImageJ.

### Statistical Analyses

Data were analyzed using GraphPad Prism 9.0. All values are mean±SEM. A nominal p<0.05 was considered statistically significant. Grubbs’ test was performed to identify and exclude outliers. Normal distribution of data was assessed using Shapiro-Wilk or D’Agostino & Pearson tests. Normally distributed data were analyzed using unpaired students *t*-tests. Non-parametric data were analyzed using Mann-Whitney *U*-tests. For correlations, Spearman analysis for non-parametric data was used. For multiple comparisons, One-way ANOVA was used, followed by Kruskal-Wallis post-tests.

## Results

### Plasma bile acids contribute to ∼80% of brain bile acid levels

BA synthesis enzymes are highly expressed in the liver, the predominant site of BA synthesis (20). However, at least part of the enzymes to synthesize primary BAs are also present in the brain (21). Thus, how much liver-derived BAs contribute to the brain BA pool remains unclear. Secondary BAs are generated from liver-derived primary BAs, solely through the action of the intestinal microbiota (2). Thus, any secondary BAs in the brain should derive from the liver generated primary BAs. We administered broad spectrum antibiotics to *wild-type* mice in order to modulate circulatory secondary BAs, and quantified these BAs in the brain. Brain secondary BAs were decreased in the antibiotics-treated mice (vehicle: 1.0±0.08, n=11; antibiotics: 0.70±0.05, n=13; fold change, p=0.003) (**Fig 1A**), suggesting that brain BAs are, in part, hepatic in origin. In addition, brain levels of the primary BA CA were decreased (vehicle: 1.0±0.13, n=11; antibiotics: 0.35±0.13, n=13; fold change, p=0.0005) and brain TCDCA was increased in the antibiotics-treated mice (vehicle: 1.0±0.23, n=11; antibiotics: 3.4±1.14, n=13; fold change, p=0.4) (**Fig 1B**).

**Figure 1.**
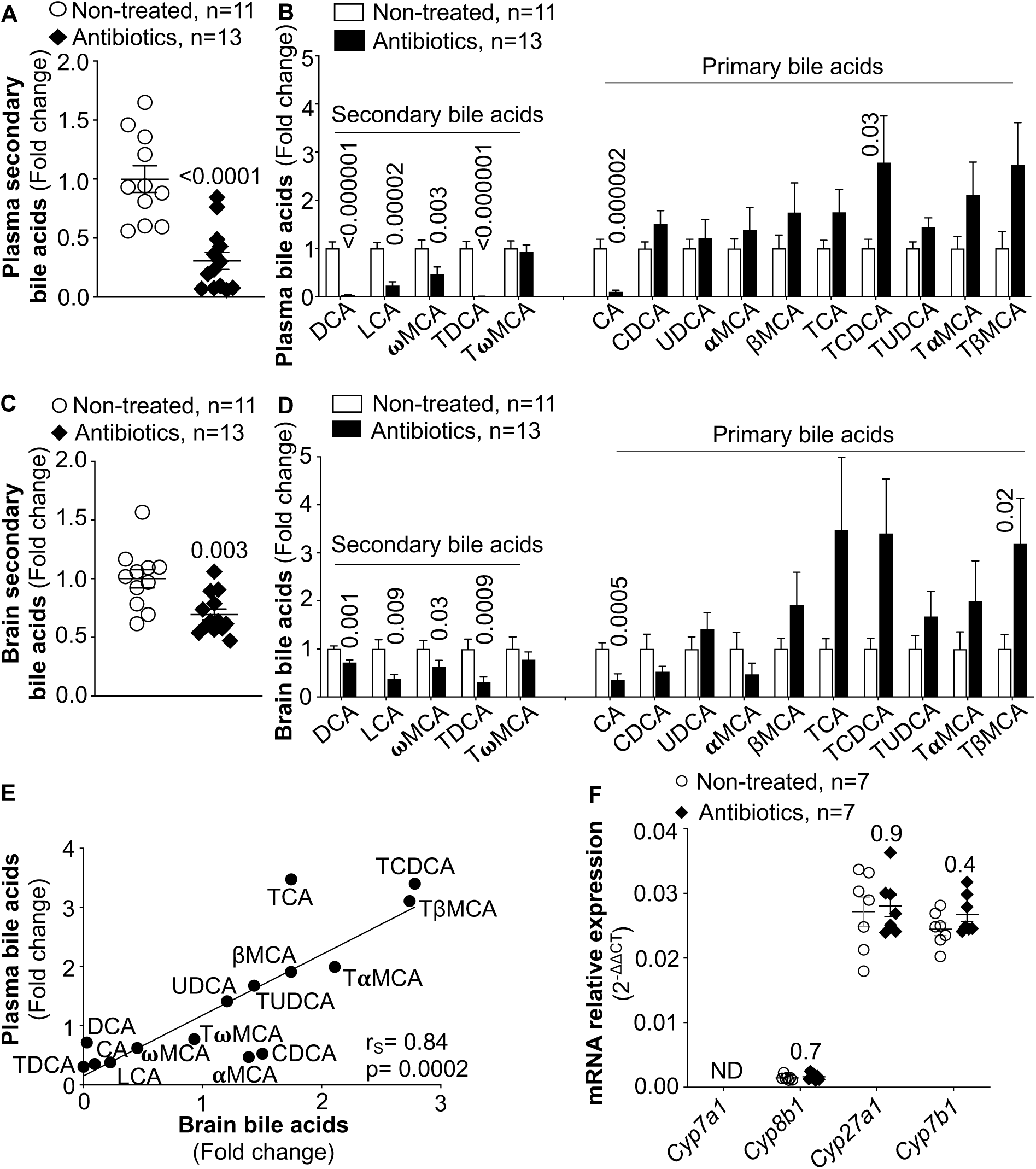
∼84% of brain bile acids are hepatic in origin. (A) Plasma secondary BAs, which are derived solely from intestinal gut bacterial function, are reduced in *wild-type* mice administered antibiotics. (B) The primary bile acid cholic acid (CA) is decreased, and CDCA and its tauro conjugate (TCDCA), as well as its derivatives TαMCA and TβMCA are increased in plasma upon antibiotic administration. (C) Brain secondary BAs are reduced in antibiotics-administered *wild-type* mice. (D) CA is decreased, and TCDCA, TαMCA and TβMCA are increased in the antibiotics-administered brains. (E) Brain BAs are strongly correlated with plasma BAs (r_s_=0.84) post antibiotic administration. (F) Antibiotic administration does not alter brain mRNA expression of the BA synthesis enzymes *Cyp8b1*, *Cyp27a1* and *Cyp7b1*. mRNA expression was normalized to Ribosomal Protein L37 (*Rpl37*). Data are shown as average±SEM, and assessed using Mann Whitney *U*-tests. ND=not detected. r_s_=Spearman correlation coefficient.

To confirm that these changes in brain secondary BAs arose due to changes in circulatory secondary BAs, we analyzed circulatory BAs in these mice. As expected, plasma secondary BAs were decreased in the antibiotics-treated mice (vehicle: 1.0±0.11, n=11; antibiotics: 0.31±0.07, n=13; fold change, p<0.0001) (**Fig 1C**). Plasma primary BAs were also altered, with decreased CA (vehicle: 1.0±0.19, n=11; antibiotics: 0.10±0.03, n=13; fold change, p=0.000002), and increased TCDCA in the antibiotics-treated mice (vehicle: 1.0±0.19, n=11; antibiotics: 2.78±0.97, n=13; fold change, p=0.03) (**Fig 1D**). Antibiotic-mediated changes in plasma BAs positively correlated with brain BA levels (r_s_=0.84, p=0.0002) (**Fig 1E**), indicating that ∼84% of brain BAs may be hepatic in origin, and suggesting that circulatory BAs are the major contributor to the brain BA pool.

To determine if altered brain BA synthesis may account for the changes in brain BAs, we quantified mRNA levels of BA synthesis enzymes in the brain. *Cyp7a1* was below detection limits, whereas *Cyp27a1*, *Cyp8b1,* and *Cyp7b1* remained unchanged in the antibiotics treated mice compared to vehicle treated mice (**Fig 1F**), suggesting unaltered brain BA synthesis.

### Brain CDCA and TUDCA levels are increased in *Cyp8b1^-/-^* mice

In order to study the effects of BAs on neuronal functions, we generated mice with deleted *Cyp8b1*, a BA synthesis enzyme critical for regulating levels of the two primary bile acids, CDCA and CA, and their derivatives (8-11). *Cyp8b1* is predominantly expressed in the liver in *wild-type* mice (20). *Cyp8b1* was not expressed in the liver of *Cyp8b1^-/-^* mice, confirming their successful generation (**Supplemental Figs 2A,B**). In plasma, *Cyp8b1^-/-^* mice showed increased CDCA (*Cyp8b1^+/+^*: 1.0±0.12, n=13; *Cyp8b1^-/-^*: 5.08±1.55, n=10; relative units, p=0.002) (**Fig 2A**) and TUDCA levels (*Cyp8b1^+/+^*: 1.0±0.13, n=13; *Cyp8b1^-/-^*: 3.71±0.47, relative units, n=10; p<0.0001) (**Fig 2B**). In line with CYP8B1’s function as a 12α-hydroxylase, plasma total 12α-hydroxylated BA levels were decreased in the *Cyp8b1^-/-^* mice (*Cyp8b1^+/+^*: 1.0±0.10, n=13; *Cyp8b1^-/-^*: 0.02±0.004, n=10; relative units, p=0.000002) (**Fig 2C**), whereas total non-12α-hydroxylated BAs were increased (*Cyp8b1^+/+^*: 1.0±0.20, n=13; *Cyp8b1^-/-^*: 1.80±0.24, n=10; relative units, p=0.008) (**Fig 2C**). These findings are consistent with previous studies in the *Cyp8b1^-/-^* mice (8-10).

**Figure 2.**
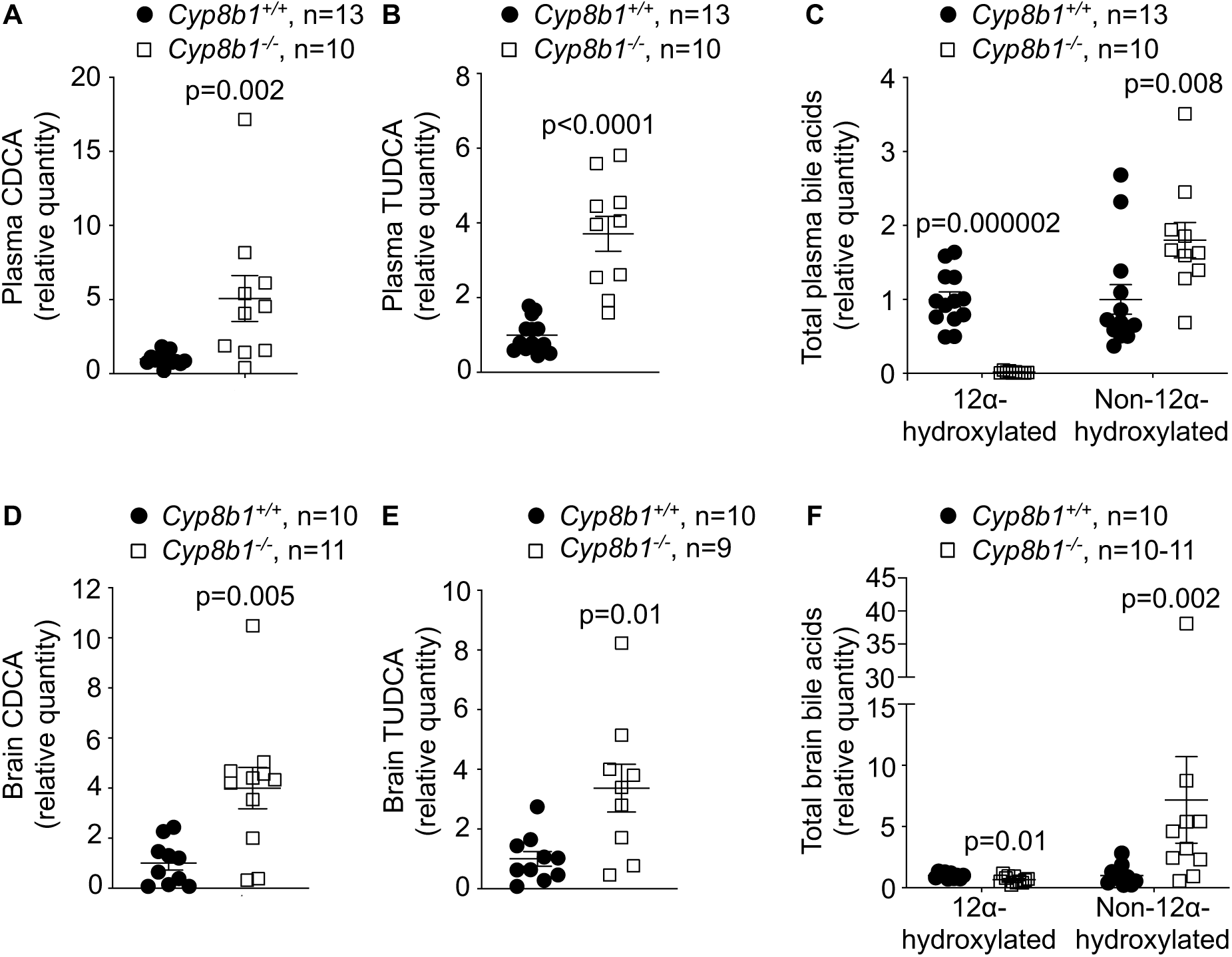
Increased brain CDCA and TUDCA levels in *Cyp8b1^-/-^* mice. Increased (A) CDCA, (B) TUDCA, and (C) non-12α-hydroxylated BAs, and decreased (C) 12α-hydroxylated BAs, in *Cyp8b1^-/-^* mouse plasma. Increased (D) CDCA, (E) TUDCA, and (F) non-12α-hydroxylated BAs, and decreased (F) 12α-hydroxylated BAs in *Cyp8b1^-/-^* brains. Data are average±SEM, and analyzed using Mann-Whitney *U*-tests.

Brain BA pool composition was assessed in *Cyp8b1^-/-^* mice perfused transcardially with PBS to remove any BA arising from brain circulation. Both CDCA (*Cyp8b1^+/+^*: 1.0±0.28, n=10; *Cyp8b1^-/-^*: 4.00±0.83, relative units, n=11; p=0.005) (**Fig 2D**), and TUDCA (*Cyp8b1^+/+^*: 1.0±0.25, n=10; *Cyp8b1^-/-^*: 3.37±0.80, relative units, n=9; p=0.01) (**Fig 2E**) were increased in *Cyp8b1^-/-^* brains. Total 12α-hydroxylated BAs were decreased (*Cyp8b1^+/+^*: 1.0±0.07, n=10; *Cyp8b1^-/-^*: 0.67±0.08, relative units, n=11; p=0.01) (**Fig 2F**), whereas total non-12α-hydroxylated BAs were increased in *Cyp8b1^-/-^* brains (*Cyp8b1^+/+^*: 1.0±0.26, n=10; *Cyp8b1^-/-^*: 7.17±3.52, relative units, n=10; p=0.002) (**Fig 2F**), mimicking their plasma BA profile.

### Reduced susceptibility to ischemic stroke in *Cyp8b1^-/-^* mice

Previous studies suggested that TUDCA administration to mouse models of neurological disorders reduced their pathological phenotypes, and CDCA administration improved neurological symptoms in patients with CTX (12-14). This raised the possibility that the brain BA composition, particularly the elevated CDCA and TUDCA in *Cyp8b1^-/-^* mice, maybe neuroprotective. Thus, we subjected the *Cyp8b1^-/-^* mice to transient middle cerebral artery occlusion (tMCAO) to induce focal cerebral ischemia, followed by reperfusion. Brain infarct area, analysed 24h after reperfusion, was decreased by 31% in *Cyp8b1^-/-^* mice (*Cyp8b1^+/+^*: 28.9±1.6, n=6; *Cyp8b1^-/-^*: 19.9±3.0, n=5; % lesion area, p=0.02) (**Fig 3A**).

**Figure 3.**
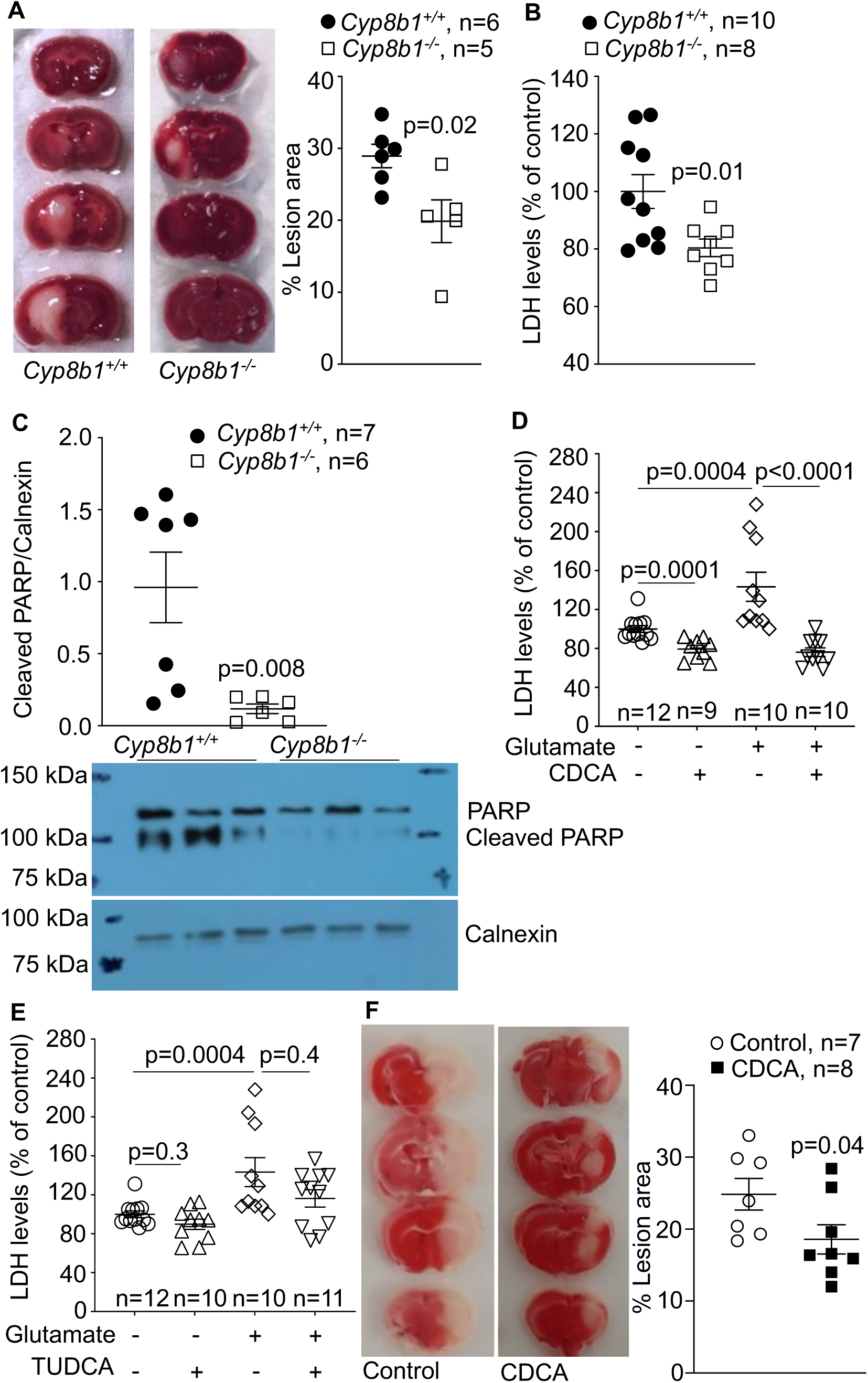
CDCA reduces focal ischemia and glutamate-induced neurotoxicity. (A) Reduced ischemic lesion area in *Cyp8b1^-/-^* mice. (B) Reduced media LDH levels, and (C) reduced 89kDa PARP-1 fragment in *Cyp8b1^-/-^* primary cortical neurons treated with glutamate. (D) CDCA reduces LDH release by glutamate-treated *wild-type* neurons. (E) TUDCA does not significantly decrease glutamate-induced LDH release in *wild-*type neurons. Experiments for (D) and (E) were performed simultaneously, therefore share controls (vehicle and 30µM glutamate). (F) Reduced ischemic lesion area in CDCA-administered *wild-type* mice. Data are average±SEM. Data in (A) and (B) were analyzed using students *t*-tests. Data in (C to F) were analyzed using Mann-Whitney *U*-tests.

A mechanism underlying neuronal death in ischemic stroke is glutamate-induced excitotoxicity (22). Since the *Cyp8b1^-/-^* mice showed reduced ischemic stroke lesion area, we investigated whether *Cyp8b1^-/-^* neurons may be protected against glutamate induced toxicity. During glutamate-mediated excitotoxicity, cytoplasmic lactate dehydrogenase (LDH) is released into the culture media due to compromised membrane integrity (23). As well, during glutamate-mediated excitotoxicity, caspase 3 cleaves poly (ADP-ribose) polymerase-1 (PARP-1), releasing an 89 kDa fragment (24). In response to glutamate, primary *Cyp8b1^-/-^* cortical neurons exhibited both reduced LDH release (*Cyp8b1^+/+^*: 100.0±5.9, n=10; *Cyp8b1^-/-^*: 80.4±3.1, n=8; % of control, p=0.01) (**Fig 3B**) and reduced cleaved PARP-1 (*Cyp8b1^+/+^*: 0.96±0.25, n=7; *Cyp8b1^-/-^*: 0.12±0.03, n=6; normalized to calnexin; p=0.008) (**Fig 3C**). Thus, the absence of *Cyp8b1* reduced the susceptibility of neurons to glutamate-induced excitotoxicity, which likely contributed to the decreased susceptibility to ischemic stroke in *Cyp8b1^-/-^* mice.

### CDCA but not TUDCA reduces glutamate-induced neurotoxicity

Both CDCA and TUDCA are increased in *Cyp8b1^-/-^* brains (**Figs 2D,E**). To determine if CDCA and/or TUDCA may contribute to neuroprotection in the *Cyp8b1^-/-^* mice, we exposed *wild-type* cortical neurons treated with the excitotoxic agent glutamate to either CDCA or TUDCA. CDCA reduced glutamate-induced LDH release from *wild-type* mouse cortical neurons (Glutamate: 143.4±14.9, n=10; CDCA+glutamate: 76.3±4.4, n=10; % of control, p<0.0001) (**Fig 3D**). Conversely, TUDCA did not significantly alter LDH release (Glutamate: 143.4±14.9, n=10; TUDCA+glutamate: 116.3±8.8, n=11; % of control; p=0.4) (**Fig 3E**). These data suggest that CDCA, but not TUDCA, mediates increased resistance of *Cyp8b1^-/-^* neurons to glutamate-induced neurotoxicity. Interestingly, CDCA alone, in the absence of glutamate, decreased LDH release (vehicle: 100.0±3.4, n=12; CDCA: 79.5±3.6, n=9; % of control; p=0.0001) (**Fig 3D**), further indicating that CDCA may promote neuronal survival.

### CDCA administration reduces ischemic stroke lesion area

To further confirm that increased brain CDCA levels are neuroprotective, we performed tMCAO on *wild-type* mice administered CDCA. Brain infarct area was decreased by 25% in the CDCA-administered *wild-type* mice (vehicle: 24.9±2.2, n=7; CDCA: 18.6±2.0, n=8; % infarct volume; p=0.04) (**Fig 3F**), suggesting that increased CDCA confers neuroprotection *in vivo*.

### CDCA decreases NMDAR-mediated excitatory postsynaptic currents (EPSCs)

Under normal conditions, glutamate acts as the main excitatory neurotransmitter (25). However, during ischemia or other excitotoxic stimuli, a substantial release of glutamate persistently activates NMDARs and α-amino-3-hydroxy-5-methyl-4-isoxazolepropionic acid (AMPA) receptors (22), leading to large Ca^2+^ influxes, resulting in excitotoxic death (25). To determine if CDCA modulates NMDAR and/or AMPAR responses, we measured NMDAR and AMPAR-mediated EPSCs in hippocampal CA1 regions of *wild-type* brain slices exposed to CDCA. The NMDAR/AMPAR-mediated EPSC ratio was decreased upon CDCA treatment (Vehicle: 1.00±0.04, n=16; 0.1µM CDCA: 0.78±0.06, n=19, p=0.003; 1µM CDCA: 0.76±0.05, n=12, p=0.007) (**Fig 4A**). AMPAR-mediated EPSCs were not altered (ANOVA p=0.7) (**Supplemental Figs 3A,B**), suggesting that CDCA decreases NMDAR-mediated EPSCs.

**Figure 4.**
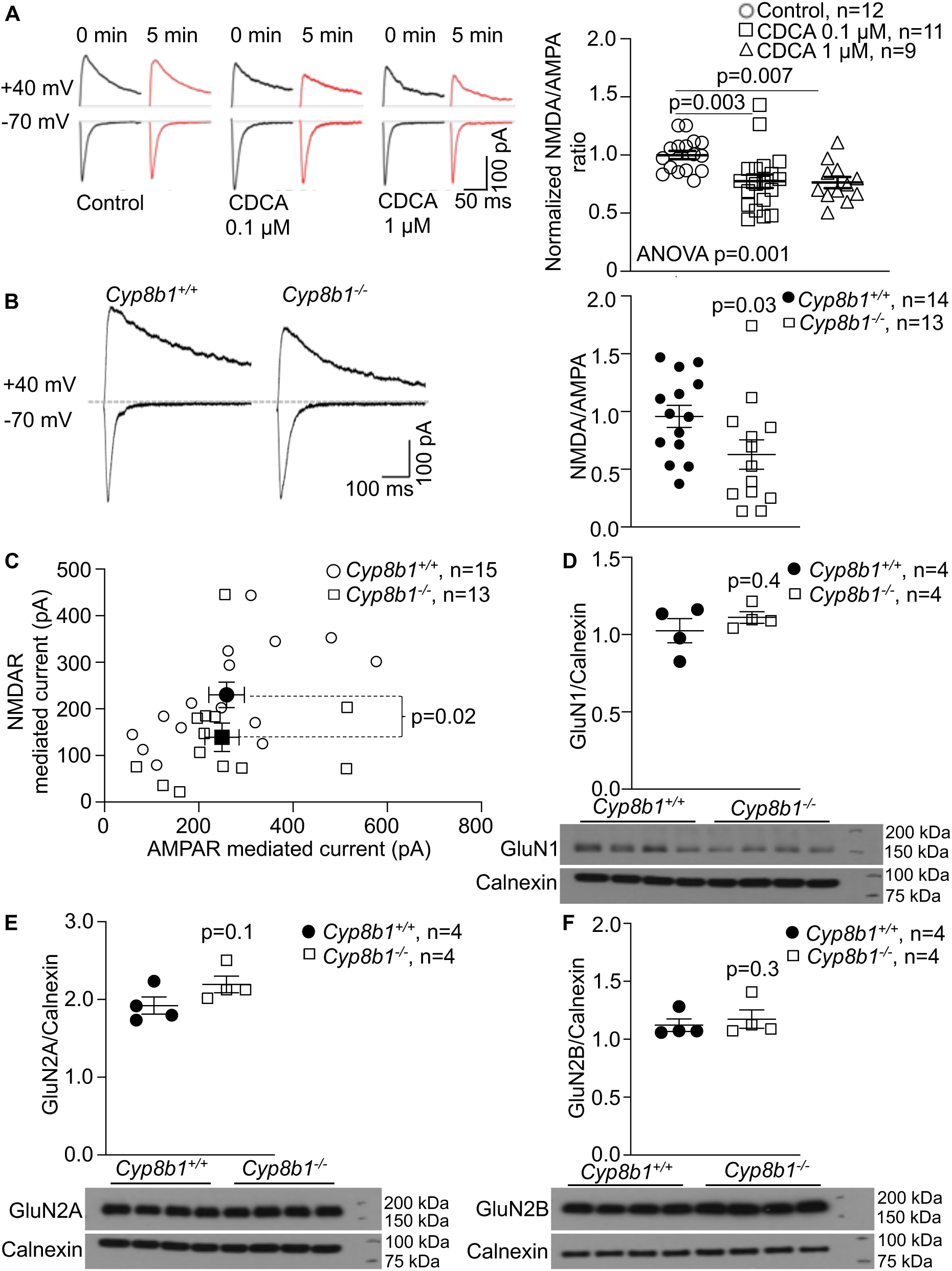
Decreased NMDAR-mediated EPSCs upon CDCA elevation. (A) Representative traces and quantitation of AMPAR-and NMDAR-mediated EPSCs in *wild-type* mouse acute brain slices showing CDCA decreases NMDAR/AMPAR ratio. (B) Representative traces of AMPAR-and NMDAR-mediated EPSCs, showing reduced NMDAR/AMPAR-mediated EPSC ratio in *Cyp8b1^-/-^* acute brain slices. (C) Reduced NMDAR-mediated EPSCs in *Cyp8b1^-/-^* mice. *Cyp8b1^+/+^* average, dark circle; *Cyp8b1^-/-^* average, dark square. (D to F) No differences in GluN1, GluN2A and GluN2B expression in *Cyp8b1^-/-^* mice. Calnexin control is shared in (D) and (E) since GluN1 and GluN2A were run on the same gel. Data are average±SEM. Data in (A) are analyzed using one-way ANOVA followed by Kruskal-Wallis post-test. (B) is analyzed using Mann-Whitney *U*-test. Data in (D to F) are analyzed using student’s *t*-tests. (B) and (C): *Cyp8b1^+/+^* n=15 from 3 mice; *Cyp8b1^-/-^* n=13 from 2 mice.

### Synaptic NMDAR-mediated EPSCs are reduced in *Cyp8b1^-/-^* mice

To investigate whether reduced NMDAR-mediated EPSCs contributed to neuroprotection in *Cyp8b1^-/-^* mice, we assessed synaptic NMDAR-mediated EPSCs in hippocampal CA1 pyramidal neurons in *Cyp8b1^-/-^* brain slices. NMDAR/AMPAR-mediated EPSC ratio was decreased in *Cyp8b1^-/-^* brain slices (*Cyp8b1^+/+^*: 0.96±0.10, n=14; *Cyp8b1^-/-^*: 0.63±0.13, n=13; p=0.03) (**Fig 4B**), due to decreased NMDAR-mediated EPSCs (*Cyp8b1^+/+^*: 230.2±27.5, n=15, *Cyp8b1^-/-^*: 138.9±30.6, n=13, pA, p=0.02) (**Fig 4C**). AMPAR-mediated EPSCs were unaltered (*Cyp8b1^+/+^*: 259.0±37.8, n=15, *Cyp8b1^-/-^*: 249.1±36.4, n=13, pA, p=0.7) (**Fig 4C**).

Reductions in NMDAR-mediated EPSCs could arise from decreased NMDAR expression levels. Most NMDARs are heterotetrametric complexes formed by two obligatory GluN1 and two GluN2 subunits, the latter consisting of either GluN2A or GluN2B (26). However, hippocampal GluN1, GluN2A and GluN2B expression was unchanged in the *Cyp8b1^-/-^* mice (**Figs 4D** to **F**). These data suggest that reduced NMDAR-mediated EPSCs likely resulted from inhibition of NMDAR-mediated EPSCs by the increased CDCA in the brains of *Cyp8b1^-/-^* mice.

### CDCA decreases GluN2B-containing NMDAR-mediated EPSCs

Excess activation of GluN2B-containing NMDARs plays a critical role in neurotoxicity in ischemic stroke (22). To determine if GluN2B-containing NMDAR activity is reduced by CDCA, we determined the decay kinetics of NMDAR-mediated EPSCs in the hippocampal CA1 region of *wild-type* brain slices exposed to CDCA (**Fig 4A**), and found them decreased (Vehicle: 202.8±24.4, n=9; 0.1 μM CDCA: 103.5±8.6, n=10; Time constant, t (ms), p=0.002), whereas AMPAR decay kinetics were not significantly altered (Vehicle: 66.6±21.4, n=9; 0.1 μM CDCA: 140.8±32.7, n=10; t (ms), p=0.1) (**Fig 5A**).

**Figure 5.**
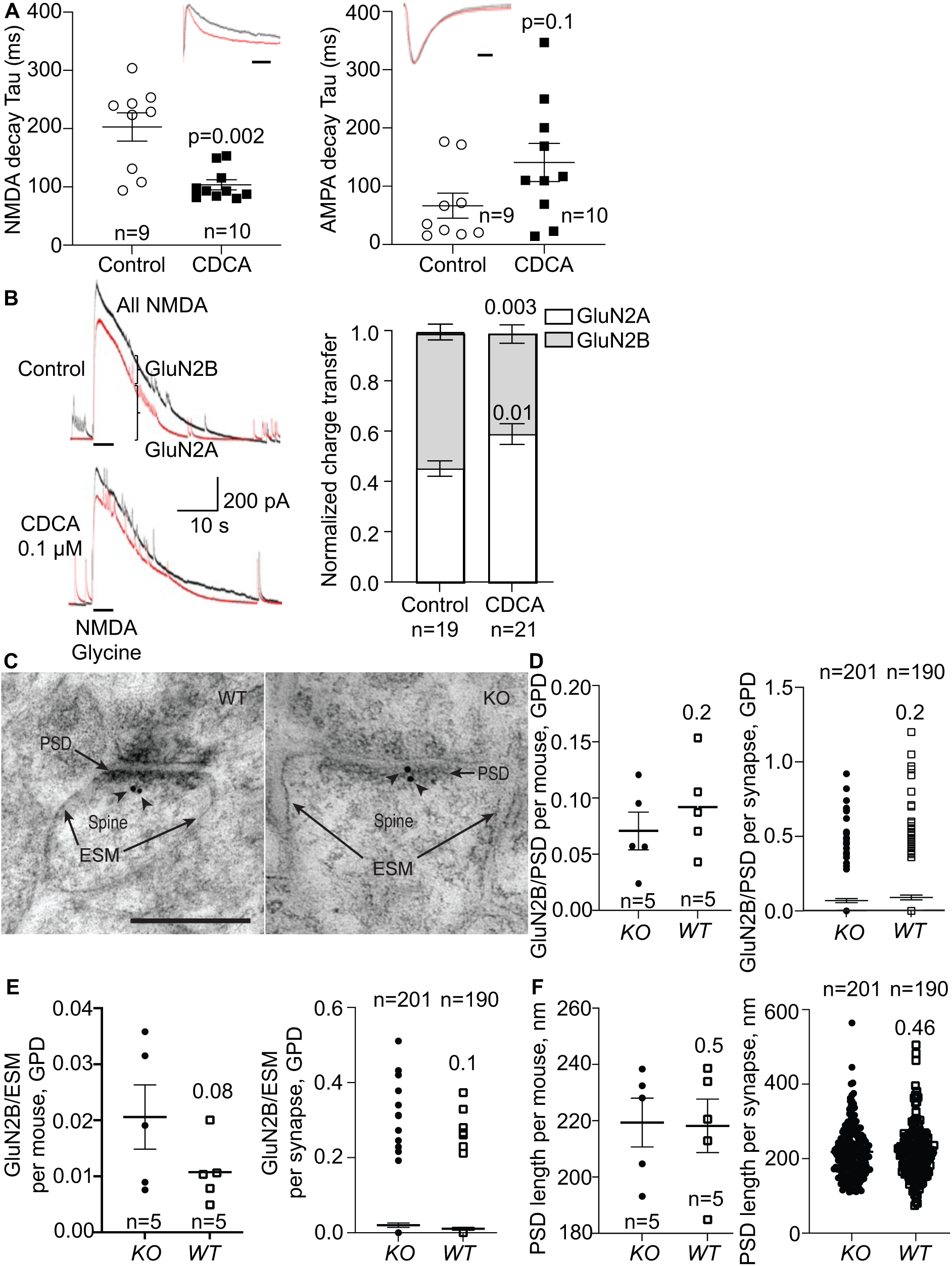
CDCA reduces NMDAR-GluN2B response, with no effects on synaptic GluN2B concentrations. (A) CDCA reduces evoked NMDAR-mediated EPSC decay time in acute hippocampal slices, but not evoked AMPAR-mediated EPSC decay time. Insets show representative traces. The CDCA amplitude (red) is scaled to control (black). Scale bar, 200 ms. (B) Representative traces of total NMDAR (black) and GluN2A (red) response from hippocampal neurons showing CDCA reduces relative GluN2B response. GluN2A response was obtained with Ro25-6981, a GluN2B-specific inhibitor, and GluN2B response was derived by subtracting GluN2A from total NMDAR response. (C) Representative post-embedding immunogold electron micrographs of GluN2B (Scale bar: 0.25µm), showing (D) unaltered gold particle density (GDP) in the post-synaptic density (PSD), per mouse, and per synapse. (E) Unaltered extra-synaptic membrane (ESM) GPD, per mouse, and per synapse. (F) Unaltered PSD length, per mouse, and per synapse. Data are average±SEM. Data in (A) and (B), and quantitation per synapse in (D to F) were analyzed using Mann-Whitney *U*-tests. Quantitation per mouse in (D to F) were analyzed using independent samples *T*-tests.

To confirm the inhibitory effect of CDCA on GluN2B-containing NMDARs, we measured NMDAR-mediated EPSCs in *wild-type* hippocampal neurons treated with CDCA and perfused with Ro 25-6981, a highly selective, activity-dependent inhibitor of GluN2B-containing NMDARs (27). A ∼30% decrease in GluN2B-mediated EPSCs was observed upon CDCA treatment (Vehicle: 0.54±0.03, n=19; 0.1 µM CDCA: 0.40±0.04, n=21; normalized charge transfer, p=0.003) (**Fig 5B**), suggesting that CDCA reduces GluN2B-containing NMDAR-mediated EPSCs. GluN2A-mediated EPSCs were not decreased by CDCA (**Fig 5B**). In conjunction with the fact that GluN2B expression is higher in the extra-synaptic region of adult mice (28), the reduced NMDAR-mediated EPSCs (**Fig 4A**) and the reduced decay kinetics of NMDAR-mediated EPSCs suggest that CDCA reduces GluN2B-mediated EPSCs in the synapse, and likely inhibits GluN2B-mediated EPSCs in the extra-synapse.

Although cellular GluN2B expression was unaltered (**Fig 4F**), reductions in GluN2B-containing NMDAR-mediated EPSCs may also arise from altered synaptic plasma membrane GluN2B distribution. Thus, we performed post-embedding immunogold electron microscopic labelling of GluN2B in the stratum radiatum of the hippocampal CA1 region in *Cyp8b1^-/-^* mice. Gold particle density (GPD, i.e., number of gold particles/µm membrane length) was quantified along the post-synaptic density (PSD), and along the extra-synaptic membrane (ESM, defined as the continuation of the synaptic plasma membrane to each side of the PSD, for a length equal to that of the PSD) (**Fig 5C**). GPD along the PSD, either per mouse (*Cyp8b1^+/+^*: 0.092±0.019; *Cyp8b1^-/-^*: 0.071±0.017, n=5 mice each, p=0.21 one-sided) or per synapse (*Cyp8b1^+/+^*: 0.091±0.016, n=201 synapses; *Cyp8b1^-/-^*: 0.069±0.014, n=190 synapses, p=0.24 one-sided, **Fig 5D**) was unchanged. As expected, GPD density along the ESM was lower than along the PSD, but not significantly different between the groups, per mouse (*Cyp8b1^+/+^*: 0.011±0.003; *Cyp8b1^-/-^*: 0.021±0.006, n=5 mice each, p=0.08 one-sided) and per synapse (*Cyp8b1^+/+^*: 0.011±0.004, n=201 synapses; *Cyp8b1^-/-^*: 0.020±0.006, n=190 synapses, p=0.14 one-sided, **Fig 5E**). PSD length was unchanged, per mouse (*Cyp8b1^+/+^*: 218.2 ±9.5; *Cyp8b1^-/-^*: 219.4±8.7, nm, n=5 mice each, p=0.5 one-sided) and per synapse (*Cyp8b1^+/+^*: 216.0±4.7, n=201 synapses; *Cyp8b1^-/-^*: 218.5±5.1, n=190 synapses, p=0.46 one-sided, **Fig 5F**). As control, no gold particles were detected along the mitochondrial membrane, showing high synaptic plasma membrane GluN2B immunogold labelling specificity. Together, these data suggest that increased CDCA in the absence of *Cyp8b1* directly reduces GluN2B activity through CDCA interaction.

## Discussion

We show here that the absence of the BA synthesis enzyme CYP8B1 in mice increases brain levels of the BA chenodeoxycholic acid (CDCA) and that CDCA reduces GluN2B-containing NMDAR over-activation that occurs during excitotoxicity, thereby decreasing neuronal death. Our findings suggest that the BA CDCA acts as a liver derived factor that modulates brain neurophysiology and signalling. However, a contribution by brain derived CDCA cannot be excluded. Secondary BAs are generated from the liver-derived primary BAs through the action of the gut microbiome. Our data show that altering the gut microbiome not only modulates secondary BA levels, but also modulates the levels of primary BAs. These findings suggest that BAs are liver generated, gut microbiome modulated signalling molecules that can act in the liver-gut-brain axis to regulate neuronal function.

CDCA reduced neuronal excitotoxicity which is a key pathological process in neurodegenerative diseases such as Huntington, Alzheimer’s, and Parkinson’s diseases (29), as well as ischemic stroke (22). Thus, our data also suggest that inhibition of CYP8B1 and the resulting increase in CDCA represents a potential therapeutic avenue for protection against excitotoxicity-mediated neuronal disorders.

Both CDCA and TUDCA are increased in *Cyp8b1^-/-^* brains. Under identical experimental conditions, our data show that CDCA significantly reduces excitotoxic neuronal death, whereas TUDCA does not. Previous studies, assessing only TUDCA, suggested that it was neuroprotective by decreasing mitochondrial depolarization, thus reducing apoptosis via a PI3K-dependent pathway (12). The presence of distinct chemical moieties affect the physical characteristics of BAs, leading to their individual activities as signalling molecules (30). Thus, it is not surprising that, compared to TUDCA, CDCA more robustly reduces glutamate-induced excitotoxicity. In line with this, of several BAs tested *in vitro*, CDCA and its conjugates were the most efficient antagonists of NMDAR (31,32).

Excitotoxicity-induced neuronal death in neurodegenerative diseases and ischemic stroke is mainly modulated by GluN2B-containing NMDARs (22,29). The selective blockade of GluN2B-containing NMDARs reduces ischemic cell death, while the selective blockade of GluN2A-containing NMDARs enhances neuronal death in ischemia (33). These findings further corroborate our data suggesting that CDCA is neuroprotective by reducing aberrant GluN2B-containing NMDAR activity.

Besides reducing aberrant NMDAR activation, CDCA may also decrease neurotoxicity via other mechanisms. CDCA is the most potent BA agonist for farnesoid X receptor (FXR) (2), which is found in the brain (4). Although the role of brain FXR is unknown, *Fxr^-/-^* mice show impaired motor coordination and cognitive dysfunction (34). As well, CDCA reduced amyloid-β-42 and improved cognition in an AD mouse model, suggested to be through FXR-mediated pathways (35). CDCA is also an agonist for Takeda G protein-coupled bile acid receptor 1 (TGR5/GPBAR1), which is also found in brain (5). Hypothalamic TGR5 regulates energy expenditure and obesity (36). The CDCA-induced antagonism of NMDAR, however, was shown to be TGR5 independent (31).

In addition to neurons, astrocytes, whose foot processes make close contact with neurons, also contain NMDARs (37). Astrocytes release gliotransmitters, including glutamate, in a feedback process regulating neuronal function (38). Overactivation of astrocytic NMDARs modulates AD, with Aβ increasing GluN2A and GluN2B expression in astrocytes, but only in the presence of neurons (39). Furthermore, NMDARs are also found in oligodendrocytes (40). Thus, detailed analyses of the effects of CDCA on NMDAR-mediated signalling in neurons, astrocytes and glia are necessary to understand the full complexity of BA-mediated regulation of NMDAR activity.

CDCA, at pharmacological concentrations, is relatively non-toxic, and its use as a therapeutic agent was approved by the US Food and Drug Administration to treat certain liver conditions and CTX (41). CDCA and UDCA, found at relatively similar amounts, are the most abundant BAs in bear bile (42). Bear bile has been used in traditional Chinese medicine for hundreds of years, with no major adverse effects. In addition, *Cyp8b1^-/-^* mice with elevated CDCA show no overt adverse phenotypes. In contrast, *Cyp8b1^-/-^* mice show improved glucose tolerance (9), decreased susceptibility to atherosclerosis (43), and decreased hepatic steatosis (10). Together, these data suggest that elevated CDCA at pharmacological levels are non-toxic, and that reduced CYP8B1 maybe protective against several disorders.

We show here that the absence of CYP8B1 results in neuroprotection, modulated in part through the inhibitory effects of CDCA on GluN2B-containing NMDA receptor activity, thus decreasing neuronal excitotoxicity. Our data suggest that increasing CDCA by inhibiting CYP8B1 might represent a therapeutic avenue to target brain disorders in which excitotoxicity plays a role. Our data also identifies a mechanism by which liver-derived signaling molecules modulate brain function.

## Acknowledgements

We thank the Singapore Bio-Imaging Consortium Nikon Imaging Centre for assistance with microscopic images.

## Supplementary Information

### Experimental procedures Antibiotic treatment

*Wild-type C57BL/6N* male mice (Taconic) between 13-15 weeks of age were submitted to a 7 day antibiotic treatment paradigm as previously described (1). Briefly, an antibiotic cocktail consisting of cephalothin (2 mg/ml, Sigma) and neomycin (2 mg/ml, Sigma), freshly prepared every day, was administered in the drinking water. After the 7 day treatment, blood was collected by orbital sinus bleeding from mice fasted for 4 hrs. Plasma was isolated and stored at −80°C until it’s use for bile acid analyses. Mice were terminally anesthetized with Ketamine (150 mg/kg) / Xylazine (10 mg/kg) solution (intra-peritoneally). Brains were dissected and snap frozen in liquid nitrogen after transcardial perfusion of the mice with 1xPBS to remove blood from the brain blood vessels and to avoid possible contamination with circulating bile acids. Brain tissue were used for BA analyses and for gene expression analysis.

### Gene expression analyses

Total RNA was extracted from brain tissue (<30 mg) using the RNeasy kit (Qiagen, Valencia, CA, USA). cDNA was generated from 2μg of RNA using random hexamers and the Superscript II First-Strand Synthesis Kit (Thermo Fisher Scientific, Waltham, Massachusetts, USA) following manufacturer’s instructions. Quantitative real-time PCR was performed using SYBR Green PCR Master Mix (Applied Biosystems, Carlsbad, CA, USA) in an ABI Quant-Studio 6 Flex Real-Time System. Reactions with 10μg cDNA were performed in technical triplicates using the mouse primers listed below. Gene expression analyses were performed by relative quantification using Ribosomal Protein L37 (*Rpl37*) as the house-keeping control, using the comparative C_t_ method (2). *Rpl37*:

Forward: 5’-GGAGTGCCAAGGCTAAGAGAC-3’, Reverse: 5’-

TCTGAATCTGCGGTAGACAATCT-3’; *Cyp7a1*: Forward: 5’-

ACGCACCTCGTGATCCTCTGGG-3’, Reverse: 5’-

GGCTGCTTTCATTGCTTCAGGGCT-3’; *Cyp8b1*: Forward: 5’-

CCTCTGGAGAAGGGTTTTGTG-3’, Reverse: 5’-GCACCGTGAAGACATCCCC-3’;

*Cyp27a1*: Forward: 5’GTTCGGTCTTGCCTGGGTCGG-3’, Reverse: 5’-

ACTTCTCCCATCCCGGGAGCC-3’; *Cyp7b1*: Forward: 5’-

CCTCTTTCCTCCACTCATACACAA-3’, Reverse: 5’-

GAACCGATCGAACCTAAATTCCTT-3’

### Generation of *Cyp8b1^-/-^* mice

All mouse protocols were conducted in accordance with the Biomedical Sciences Institute, Singapore, institutional animal care committee guidelines. Sperm from *Cyp8b1^tm1(KOMP)Vlcg^* were obtained from the University of California, Davis, Knockout Mouse Project (KOMP) repository, and *Cyp8b1^+/-^* mice were generated by *in vitro* fertilization on the *C57BL/6N* background. The mice were intercrossed to generate *Cyp8b1^-/-^* and *wild-type* littermates, and all pups were segregated by genotype at weaning to avoid oral-fecal contamination of BAs. To confirm the deletion of *Cyp8b1*, expression of *Cyp8b1* mRNA was assessed in the liver, the major site of *Cyp8b1* expression. Experiments utilized male mice, housed in 12-hour light/dark cycles and with *ad libitum* access to water and food (Altromin 1324).

### Neuronal cell culture and cytotoxicity assay

Mixed cortical and glial cell cultures were prepared as previously described (3). Cortices were dissected from post-natal day 0 pups in Ca^2+^/Mg^2+^-free Hank’s balanced salt solution supplemented with 0.11 mg/mL sodium pyruvate, 0.1% (w/v) D-glucose and 10 mM HEPES. Tissues were dissociated in Tryple^TM^ Express (Gibco) for 10 min at room temperature. Cortices were washed twice with the plating medium (MEM Eagle’s media with Earle’s BSS (Invitrogen) supplemented with 10% FBS, 0.45% D-glucose, 0.1 mM sodium pyruvate, 0.2 mM glutamine, 10 mM HEPES, penicillin (100 U/ml) and streptomycin (100 µg/ml)) and dissociated by pipetting up and down with a 5 mL pipet. The cell suspension rested for 10 min at room temperature. The supernatant (avoiding the debris) was transferred into a new tube and homogenized by gentle pipetting. Neurons were plated on poly-L-lysine (Sigma) coated plates in the plating medium at a density of 5x10^5^ cells/cm^2^. 4 to 6 hours after plating, the plating medium was replaced by Neurobasal medium (Invitrogen) supplemented with 2% B-27 supplement (Invitrogen), 0.2 mM glutamine, penicillin (100 U/ml) and streptomycin (100 µg/ml). A day after plating, media was replaced with fresh media, and subsequently, half the media was replaced every 3-4 days. After 14 days in culture, cells were incubated either with 30 µM L-glutamate (Sigma), 30 µM CDCA (Sigma) or 30 µM CDCA plus 30 µM L-glutamate, 100 µM TUDCA or 100 µM TUDCA plus 30 µM L-glutamate. The cells were pre-incubated for 3 hrs with CDCA or TUDCA before glutamate was added. Cytotoxicity was evaluated in the culture media after 24 hour treatment, using Pierce LDH Cytotoxicity Assay Kit (Thermo Scientific) following manufacturer’s protocol. Hippocampi were isolated from postnatal day 0 pups in ice cold Ca^2+^/Mg^2+^-free Hank’s balanced salt solution. Tissues were washed thrice with dissociation medium before dissociation in 0.25% Trypsin (Gibco) for 10 minutes at 37°C, prior to pipetting up and down with a 5 ml serological pipet in the presence of DNase I. Trypsin was deactivated with an equal amount of fetal bovine serum and the cell suspension was passed through a 70 μm cell strainer. The resulting cell suspension was spun at 120 g for 5 mins. The cell pellet was gently pipetted with a 5 mL pipet and plated on poly-L-lysine (Sigma) coated 12 mm coverslips in plating medium containing DMEM, High Glucose (Gibco), 10 mM HEPES, 0.1 mM non-essential amino acids (Invitrogen), 2 mM GlutaMAX (Gibco) 1 mM Sodium Pyruvate, 10% fetal bovine serum (Gibco) and penicillin (100 U/ml) and streptomycin (100 μg/ml). The plating medium was fully replaced by Neurobasal medium (Invitrogen) supplemented with 2% B-27 supplement (Invitrogen) the next day. Subsequently, half the media was replaced every 3-4 days.

### Bile acid measurements

BAs were analyzed in plasma and brains from mice that were fasted for 4 hours. For plasma isolation, blood was drawn from the saphenous vein. Mice were then anaesthetized with ketamine (150 mg/kg)/xylazine (10 mg/kg) and transcardially perfused with 1 x phosphate-buffered saline (PBS). Brains were dissected and snap frozen. BAs were analyzed by ultra-performance liquid chromatography electrospray ionization-tandem mass spectrometry in multiple reactions monitoring (MRM) mode (UPLC-MRM MS, Genome BC Proteomics Center, University of Victoria, Canada) as follows:

Plasma: 30 µL plasma, 30 µL internal standards and 240 µL of MeOH-acetonitrile (1:1) were mixed, vortexed, sonicated in an ice-water bath, and centrifuged. The supernatant was subjected to reversed-phase solid phase extraction (SPE) using Strata-X SPE cartridges (60 mg/1mL). BAs were eluted with 1 mL x 2 of methanol-acetonitrile (1:1). The fractions were dried in a nitrogen evaporator, dissolved in 120 µL of 50% methanol, and 20 µL was injected for UPLC-MRM/MS.

Brain: Brains were weighed and homogenized. After centrifugation, 14 µL/mg tissue of acetonitrile was added and samples homogenized, and sonicated. After centrifugation, 1 mL of supernatant was mixed with 25 µL internal standard solution and subjected to phospholipid depletion-solid phase extraction (PD-SPE). Flow-through fractions were pooled, dried in a nitrogen evaporator, reconstituted in 50 µL of 50% methanol, and 20 µL was injected for UPLC-MRM/MS. Samples were analyzed in an Agilent 1290-UHPLC coupled to a Sciex-4000 QTRAP mass spectrometer which was operated in multiple-reaction monitoring (MRM) mode with negative-ion detection. A Waters BEH-C18 column (2.1 mm I.D. x 15 cm, 1.7 µm) was used for LC separation with a mobile phase of (A) 0.01% formic acid in water and (B) 0.01% formic acid in acetonitrile for binary-solvent gradient elution. Standard mixtures were dissolved in 50% methanol at 10 nmol/mL each, further diluted 1:4 (v/v), and 50 µL of each was mixed with 50 µL of an internal standard containing 14D-labeled BA analogues. 20 µL of each was injected for UPLC-(-)ESI-MRM/MS. Linear calibration curves were constructed using analyte-to-internal standard peak area ratios (As/Ai) versus molar concentrations (nmol/mL) of each BA. Data was normalized to average of *Cyp8b1^+/+^*.

### Transient middle cerebral artery occlusion (tMCAO) in *Cyp8b1^-/-^* mice

tMCAO was carried out as previously described (4). Male mice (4.5 - 6 months) were anesthetized with isoflurane. An incision was made in the neck of the mice and the left common carotid artery (LCCA) was isolated and ligated using a 1.5 cm long 0/7 silk suture. The left external carotid artery (LECA) was separated and a second ligature was made. The left internal carotid artery (LICA) was isolated and ligated with 7/0 silk suture. A small incision was made in the LCCA before it bifurcates and a 0.21 mm MCAO silicone-rubber coated nylon monofilament (Doccol Corporation, USA) was introduced into the LICA for 9-10 mm, to occlude the origin of the left middle cerebral artery at the circle of Willis. The ligature on the LICA was closed to fix the filament in position. After 1 hour of artery occlusion, the nylon monofilament and common carotid artery ligature were removed to initiate brain reperfusion. The retractors were removed and the muscle layer and the two fat bodies returned to the physiological position. The skin was closed with three separated metallic clips and the mouse placed on its left side under the infrared light until awake. Mice were sacrificed 24 hours post-MCAO, and brains isolated and sectioned in 2 mm coronal sections. Lesion area was identified by incubating brain sections with 2% 2,3,5-triphenyltetrazolium (TTC, Sigma) in PBS at 37°C for 5 minutes, followed by washing. Images were captured and infarct area quantified using Image-J. Data are expressed as % of infarct area/total brain area.

### Transient middle cerebral artery occlusion in CDCA administered *wild-type* mice

Male 12 week-old *C57BL6/N* mice were administered 1% CDCA in chow diet for 7 days. Transient middle cerebral artery occlusion was then carried out. Male mice (12 weeks) were anesthetized with Zoletil/xylazine. An incision was made in the neck of the mice and the left common carotid artery (LCCA) was isolated and ligated using a 1.5 cm long 0/6 silk suture. The left external carotid artery (LECA) was separated and a second ligature was made. The left internal carotid artery (LICA) was isolated and clamped with a microvessel clamp. A small incision was made in the LCCA before it bifurcates and a silicone-rubber coated nylon monofilament size 6-0, length 20mm; diameter with coating 0.21±0.02 mm; coating length 1-2 mm (Doccol Corporation, USA) was introduced into the LICA for 9-10 mm, to occlude the origin of the left middle cerebral artery at the circle of Willis. The ligature on the LICA was closed to fix the filament in position. After 1 hour of artery occlusion, the nylon monofilament and common carotid artery ligature were removed to initiate brain reperfusion. The retractors were removed and the muscle layer and the two fat bodies returned to the physiological position. The skin was closed with surgical glue and the mouse placed on its left side on a heating pad until awake. Mice were sacrificed 24 hours post-MCAO, and brains isolated and sectioned in 2 mm coronal sections. Lesion area was identified by incubating brain sections with 2% 2,3,5-triphenyltetrazolium (TTC, Sigma) in PBS at 37°C for 5 minutes, followed by washing. Images were captured and infarct area quantified using Image-J. Data are expressed as % of infarct area/total brain area.

### Preparation of brain slices and whole cell patch

Mice (3-4 week old males) were euthanized by cervical dislocation, brains immediately removed, transferred to ice-cold oxygenated artificial cerebrospinal fluid (aCSF) containing 124 NaCl, 2.5 KCl, 1.2 NaH_2_PO_4_, 24NaHCO_3_, 5 HEPES, 12.5 Glucose, 2 CaCl_2_, 2 MgSO_4_ (in mM, pH 7.3), and sliced horizontally at 350 µm thickness using a VT-1000 vibratome (Leica, Germany). After incubation in aCSF at 37°C for 30 min, brain slices containing the hippocampus were transferred to a recording chamber containing oxygenated aCSF with 50 µM picrotoxin. Recording pipettes were pulled from borosilicate glass pipettes using P-1000 (Sutter Instruments, USA), and pipettes with 4-7 MΩ of resistance were filled with internal solution containing 115 Cs-Gluconate, 20 CsCl, 10 HEPES, 2.5 MgCl_2_, 4 Na-ATP, 0.4 Na-GTP, 10 Na phosphocreatine, 0.6 EGTA and 5 QX-314 (in mM, pH 7.2) and used for whole-cell patching. AMPAR and NMDAR-mediated EPSCs of the CA1 pyramidal neurons were evoked by stimulation of Schaffer collateral fibers in the stratum radiatum of CA2 using tungsten stimulating electrode and electrical stimulator (A-M Systems, USA). NMDAR-mediated EPSC was measured at +40 mV of holding potential and the amplitude of NMDAR-mediated EPSC was considered 50 ms after stimulation. AMPAR-mediated EPSC was measured at −70 mV of holding potential and the amplitude of AMPAR-mediated EPSC was measured at peak response. Signals were amplified by Multiclamp-700A amplifier (Axon Instruments, USA) and digitized on Digidata-1550B (Axon Instruments, USA) and pCLAMP v10.

### Functional inhibition of NMDAR-mediated excitatory postsynaptic currents (EPSCs) by extra-cellular CDCA

Whole-cell patch was conducted in hippocampal CA3-CA1 synaptic circuits from brain slices from WT mice, with cesium-based internal solution. AMPAR and NMDAR-mediated EPSCs were measured at −70 mV and +40 mV of holding membrane potentials in CA1 pyramidal neurons with electric stimulation. In each cell, AMPAR-and NMDAR-mediated EPSCs were measured for 5 minutes before (baseline) and for 5 min after perfusion with 0.1 µM or 10 µM of CDCA dissolved in aCSF. To avoid effects from residual binding of CDCA, we collected results from only a single cell per brain slice.

### NMDA/AMPA ratio, input/output curves of AMPAR current

The NMDA/AMPA ratio and input/output curves of AMPAR current were measured at the CA1 pyramidal neurons after electrical stimulus on Schaffer collateral evoked with the Model 2100 Isolated Pulse Stimulator (A-M system). Before measurement of the NMDA/AMPA ratio and input/output curves, the electrical stimulus was evoked for 5-10 minutes to synchronize AMPAR and NMDAR mediated currents. Stimulation pulses were delivered every 30 seconds. 0.1 μM or 1 µM CDCA and 50 μM picrotoxin were included in the aCSF and directly perfused onto the hippocampal brain slices for 5 min. AMPAR and NMDAR mediated excitatory currents were measured at −70 mV and +40 mV, respectively. The input/output curves were obtained with measurements of AMPAR currents at 0.5, 1, 2, 4, 6, 8, 10 mA of electrical stimulation. AMPAR current was measured at the peak of response, and NMDA current at 50 ms. The NMDA/AMPA ratio was calculated by dividing the amplitude of NMDAR current by amplitude of AMPAR current.

### Western immunoblotting

Hippocampal protein lysates from 4-5 month old mice were obtained by homogenization and sonication in buffer containing 50 mM Tris, pH 7.5, 150 mM NaCl, 1 mM EGTA, 1% Triton X-100, 100 g/ml phenylmethylsulfonyl fluoride, and protease inhibitors (Roche Diagnostics, Germany). Primary neuron protein lysates were obtained by homogenization and sonication in RIPA buffer (Sigma) supplemented with protease inhibitors (Roche Diagnostics). Lysates were centrifuged at 15,000 x g, and 30 µg protein samples were resolved on 7.5% SDS-PAGE gels (TGX™ FastCast™ Acrylamide Kit, BioRad), and transferred onto nitrocellulose membranes. Membranes were blocked with 5% non-fat milk for 1h and incubated overnight at 4°C with primary antibodies against GluN1, GluN2A, GluN2B and PARP (1:1000, Cell Signalling Technologies, USA) in 5% BSA, or calnexin (1:5000, Sigma-Aldrich) in 5% non-fat milk. Membranes were incubated with HRP-conjugated secondary anti-rabbit antibody (Cell Signalling Technologies, 1:2000, in 5% BSA for GluN1, GluN2A, GluN2B and PARP, and 1:5000 in 5% non-fat milk for calnexin). Signals were detected using Clarity Western ECL (BioRad). Relative densities of GluN1, GluN2A and GluN2B vs. calnexin were calculated from the percent area under the curve generated by ImageJ.

### GluN2B blocker puffing experiments on hippocampal neurons

NMDAR current was measured from DIV 15-20 primary hippocampal neurons through whole-cell patch-clamp which was performed with the recording pipette filled with internal solution containing CH_3_CsO_3_S 135, NaCl 8, EGTA 0.3, HEPES 10, Na-phosphocreatine 7, Mg-ATP 4, Na-GTP 0.3, 5 QX-314 (in mM). The recording was performed in a chamber perfused with an external cellular solution containing (in mM): NaCl 124, KCl 2.5, NaH_2_PO_4_ 1.2, NaHCO_3_ 24, HEPES 5, glucose 12.5, MgSO_4_ 1 and CaCl_2_ 2. 50 μM picrotoxin and 1 μM Ro 25-6981 hydrochloride hydrate (Sigma Aldrich) were added into the external solution to block GABA dependent responses from hippocampal inhibitory neurons and GluN2B-specific NMDAR response, respectively. 100 µM NMDA with 10 µM glycine (Sigma Aldrich) was dissolved in external cellular solution and perfused into the chamber using a Picospritzer III microinjector (Parken Instrumentation) for 5 seconds at 50 µm distance away from the recording cell. Currents were measured for 60 seconds at +40 mV with voltage clamp. Each cell was recorded in triplicate, with at least 30 second intervals between each sweep. All recording pipettes were pulled from borosilicate glass pipettes using P-1000 (Sutter instrument, USA), and pipettes with 4-7 MΩ of resistance were used for whole cell patching. The recording signals were amplified using MultiClamp 700b and digitized with Digidata 1550b (Molecular Devices) and analyses were performed using Axograph X (Axogrpah Scientific) or Clampfit 11.1 (Axon Instruments).

### Electron microscopy analyses of GluN2B synaptic/extra-synaptic localization

Male *Cyp8b1^-/-^* and *wild-type* littermate mice were deeply anaesthetized with ZRF cocktail (Zoletil 250 mg, Rompum 20 mg/ml, Fentanyl 50 mg/ml is sterile isotone saline) through intraperitoneal injections immediately before being sacrificed by transcardial perfusion with 10–15 sec flush of 4% Dextran-T70 in sodium phosphate buffer (pH 7.4), followed by fixation with 0.5 L of a mixture of 4% formaldehyde and 0.1% glutaraldehyde in the same buffer. The brains were dissected out and stored in a mixture of 0.4% formaldehyde and 0.01% glutaraldehyde in sodium phosphate buffer (pH 7.4).

Tissue pieces (0.5 × 1.0 mm) of the dorsal hippocampal CA1 region were freeze substituted, sectioned, and immunolabeled as described previously (5). The tissue blocks were cryoprotected in increasing concentrations of glycerol (30 min in 10%, 30 min in 20%, and overnight in 30% at 4°C) in 0.1 M phosphate buffer and then frozen in a cryofixation unit (Reichert KF80, Vienna, Austria) filled with propane which was cooled down by liquid nitrogen. The tissue pieces were placed in Reichert capsules in a flow through chamber filled with 1.5% uranyl acetate diluted in anhydrous methanol in a precooled chamber (−90°C) in a Reichert automatic freeze substitution unit (AFS) (Leica, Germany). Following 30 h at −90°C, the temperature was raised by 4°C increments per hour from −90 to −45°C. The tissue pieces were rinsed with anhydrous methanol and infiltrated with Lowicryl HM20 resin (Polysciences Inc., Warrington, PA; Cat#15924) at a temperature of −45°C. Infiltration in 1:1 and 2:1 methanol to pure Lowicryl lasted for 2 h each, followed by 2 h and then overnight in pure Lowicryl. The Reichert capsules were moved to Lowicryl-filled gelatin capsules and then transferred to another container filled with ethanol. The resin/tissue was polymerized with UV-light for 24 h at −45°C. The temperature was increased by 5°C increments to 0°C where it was polymerized for further 35 h. Ultrathin sections (90 nm) were cut with a diamond knife (Diatome ultra 45°, Diatome) on an ultramicrotome (Reichert Ultracut S-2.GA-E-12/92, Leica Microsystems, Germany) and placed on coated (Coat-Quick “G”) nickel grids (Electron Microscopy Sciences, G300-Ni).

Postembedding immunogold labeling was carried out by incubating the sections in TBST (Tris buffered saline with Tween20, pH 7.4) containing 50 mM glycine (10 min) and in TBST containing 5% normal goat serum (GSA) (10 min) to neutralize free aldehyde groups and blocking nonspecific antibody binding sites respectively. The sections were incubated with anti-GluN2B (1:50 in 5% NGS in TBS with 0.1% Triton X-100, for 2 h at room temperature. The sections were rinsed and immersed in TBST (10 min), before they were incubated in goat anti-rabbit IgG colloidal gold-secondary antibody 1:20, in 5% NGS and 1 mg/mL PEG in TBST for 90 min. The sections were then rinsed with distilled water and post-stained with 2% uranyl acetate (90 s) and 0.3% lead citrate (90 s). Uranyl acetate and lead citrate were removed with distilled water, and sections were left to dry completely before use in the electron microscope. All sections were labeled blinded on a grid support plate at room temperature.

Electron micrographs were acquired using a transmission electron microscope (Tecnai G2 Spirit, FEI Company) operating at high tension (80 kV) with a 43,000× magnification. Electron micrographs were obtained at random from the middle layer of the stratum radiatum of the dorsal CA1 region of the hippocampus. Immunolabeling was quantified as number of gold particles/μm of membrane length in asymmetric synapses. Asymmetric synapses in this location are well-defined glutamatergic synapses, where the presynaptic element is the nerve ending of the Schaffer collateral neurons they originate from. Excitatory synapses were identified by the presence of two closely aligned membranes with a synaptic cleft, a prominent PSD and circular synaptic membrane-orientated vesicles at the presynaptic side. Only synaptic profiles with clearly visible synaptic membranes and a PSD were selected for quantitative analysis. The images were quantified by ImageJ. For the quantification of PSD, the plugin “distance to path” was used in ImageJ (6), to study the linear density of gold particles along the PSD. The segmented line tool was used to outline the region of interest, PSD, and the gold particles were marked with the multipoint tool. This was done to every synapse that was included from the micrographs. The outlined path and the points were then saved with a corresponding dtp file containing the coordinate information. All micrographs with the coordinates of the PSD and the gold particles were then run with the python component of the software tool to get the data from each synapse. The spatial resolution is the resolution for the post-embedding immunogold technique and must be determined to distinguish whether gold particles are more likely to being attached to the postsynaptic membrane. Theoretically, the resolution of this technique depends on the distance between the epitope and the center of the gold particle. This distance represents the radius of the gold particle and the diameter of both antibodies (primary and secondary). 12 nm gold particles were used, and with a radius of 6 nm in addition to the length of both the primary and the secondary antibody, 8 nm each, the spatial resolution was set to 22 nm in python. The program produced information of all gold particles within 22 nm of distance to the PSD line, and the length of the PSD. The output was produced in excel-files. The column used from the excel-file were “path length” and “particles within 22 nm of path”. With this information, linear gold particle density was calculated for each synapse. The data were then converted from nm to μm. The gold particle density (GPD) was quantified as number of gold particles per μm along the plasma membrane. After quantification, the data were imported into SPSS for further statistical analysis. Statistical analysis was performed using IBM SPSS Statistics version 27. The Shapiro-Wilk normality test was performed to determine normal data distribution. Normally distributed data were analyzed with an Independent Samples t-Test to compare the two groups. Non-normally distributed data were analyzed with Mann-Whitney U tests. Statistical significance level was set to p<0.05. Normally distributed data was presented as mean±SD while non-normally distributed data was presented as median±SEM.

---

**Supplemental Table 1:**
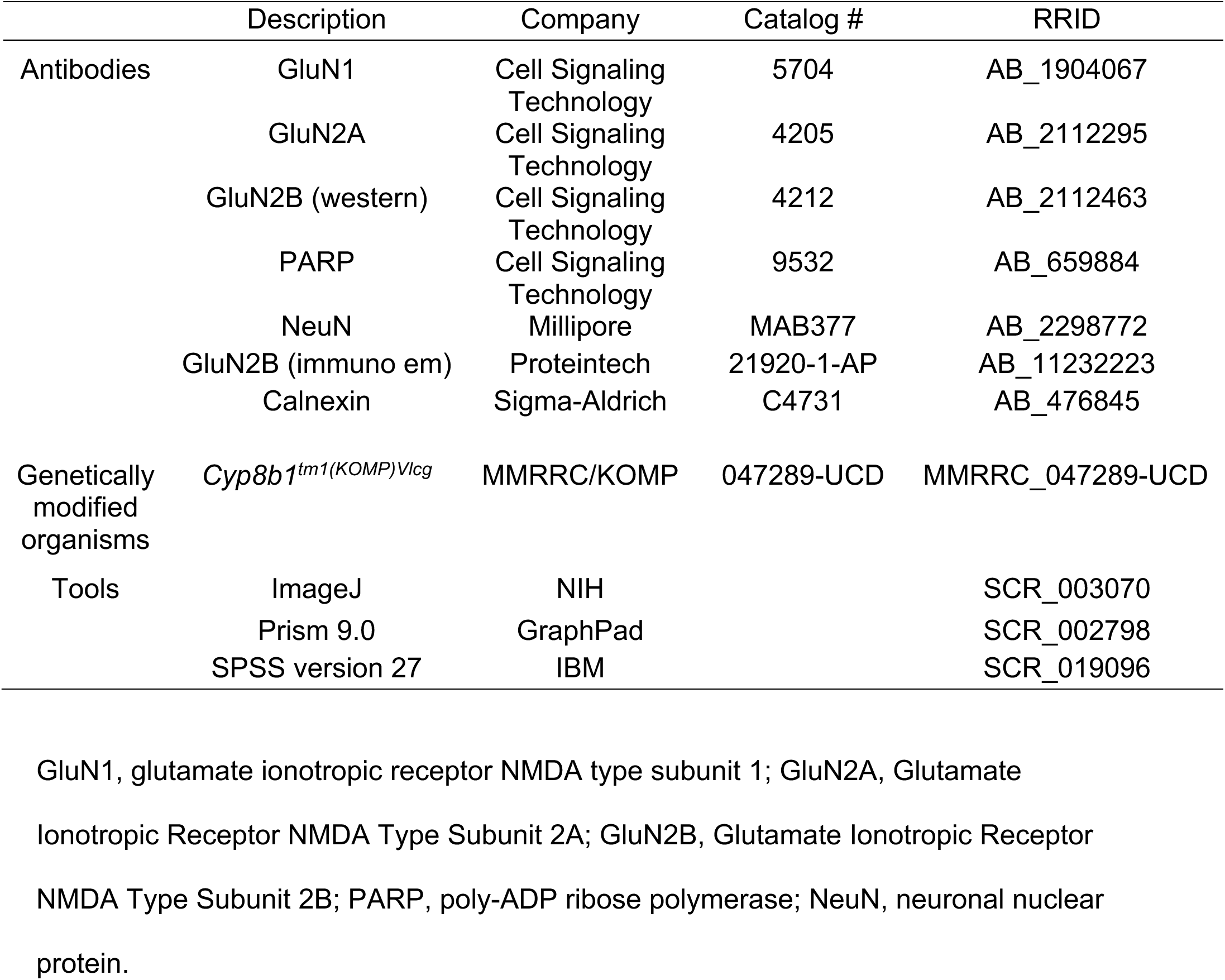
Key reagents utilized in this study.

**Supplemental Figure 1.**
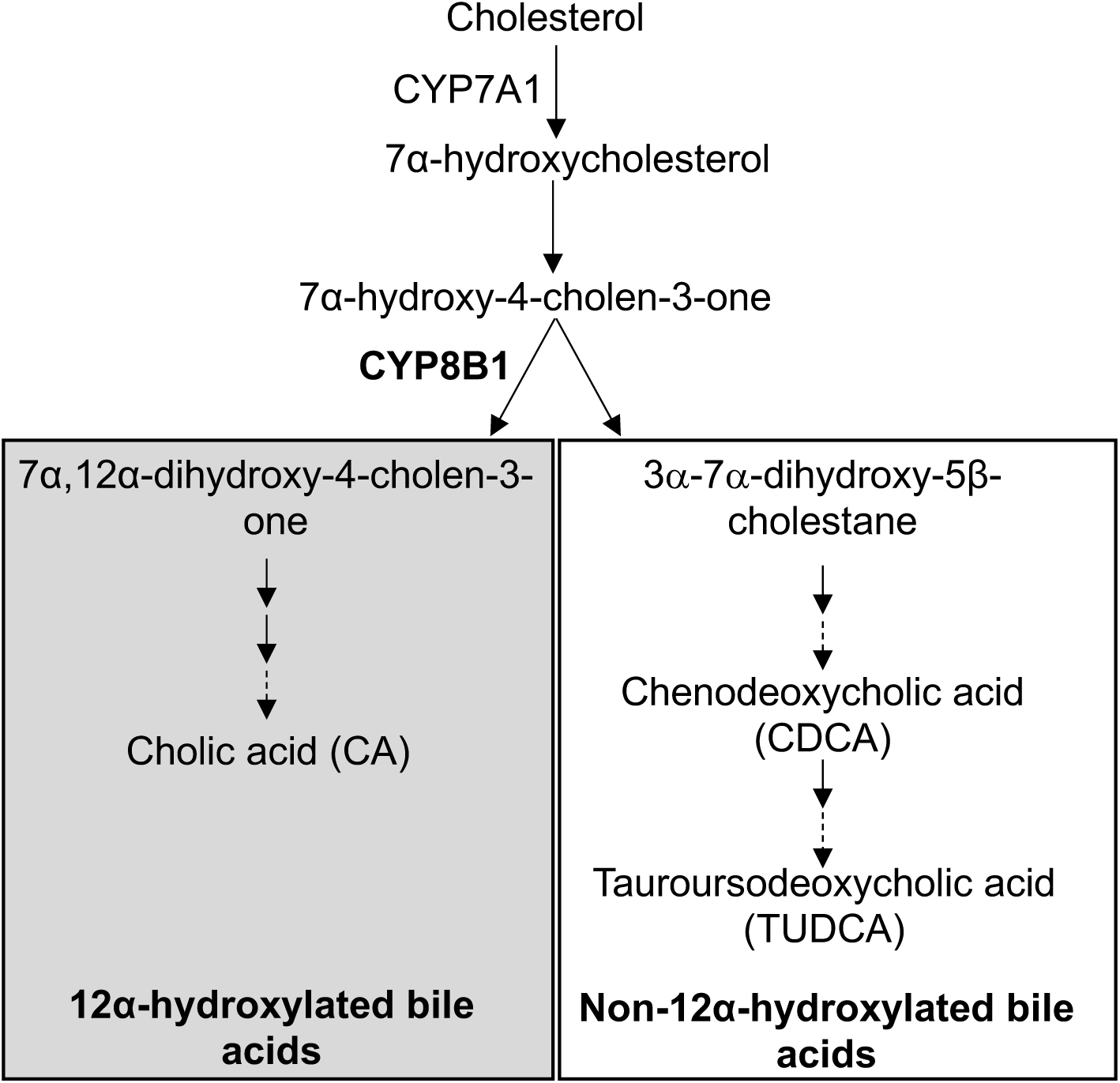
Schematic of the neutral BA synthesis pathway. The neutral bile acid synthesis pathway is the major pathway of BA synthesis and occurs in the liver. Cytochrome P450 Family 8 Subfamily B Member 1 (CYP8B1) generates the primary BA cholic acid (CA). In the absence of CYP8B1, CA is reduced, and the primary BA chenodeoxycholic acid (CDCA) is increased. Before secretion from the liver, primary BAs are conjugated with either glycine or taurine.

**Supplemental Figure 2:**
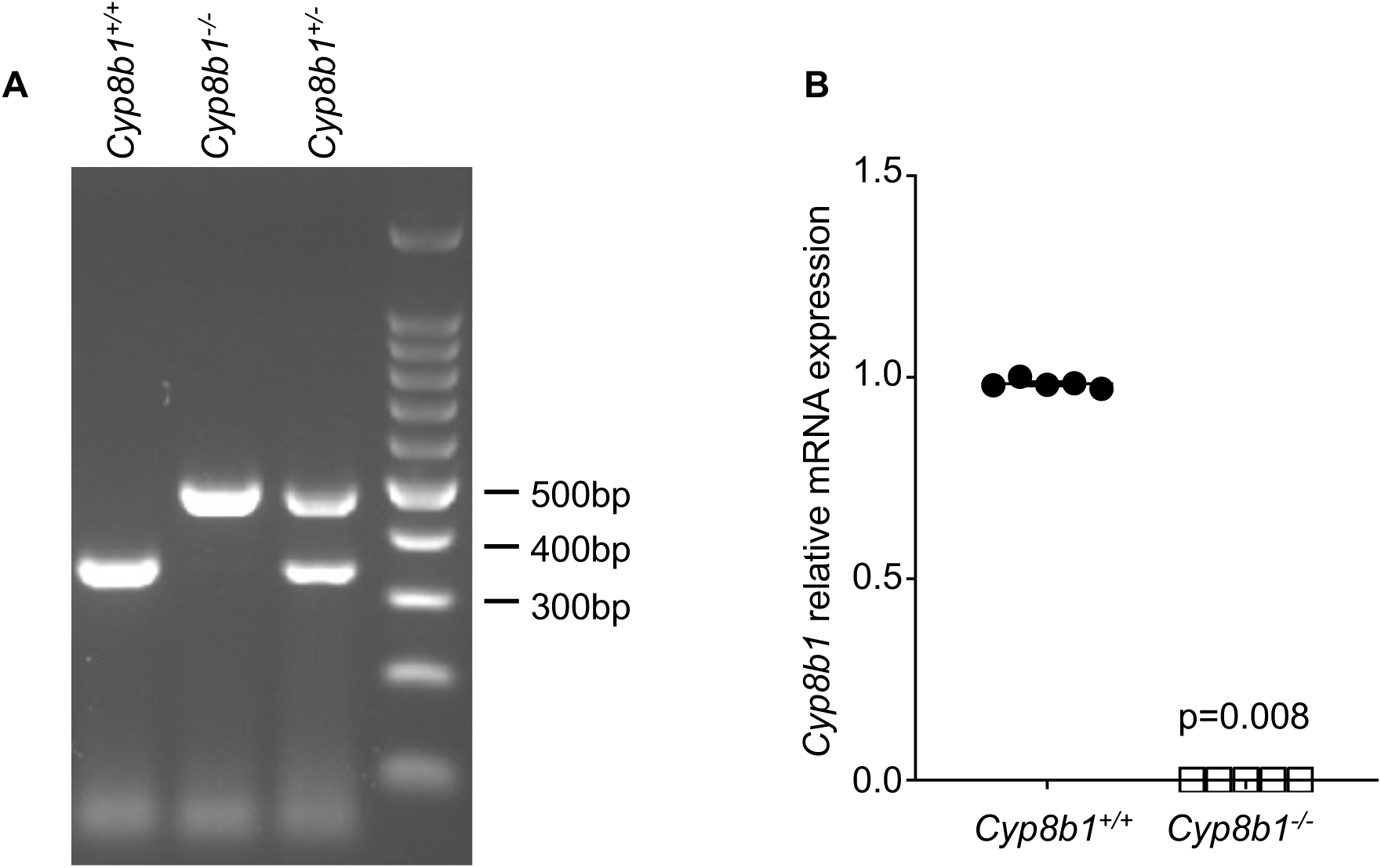
Generation of *Cy8b1^-/-^* mice. (A) Genotyping PCR showing *Cy8b1^+/+^*, *Cy8b1^+/-^*, and *Cy8b1^-/-^* mice, and (B) quantitative RT-PCR of the *Cyp8b1* transcript from the liver of *Cy8b1^+/+^* and *Cy8b1^-/-^* mice, showing no *Cyp8b1* expression in the *Cy8b1^-/-^* mice. Data is shown as average ± SEM of n=5 mice and were normalized to Ribosomal Protein L37 (*Rpl37*). Data were analyzed using the Mann-Whitney *U*-test.

**Supplemental Figure 3:**
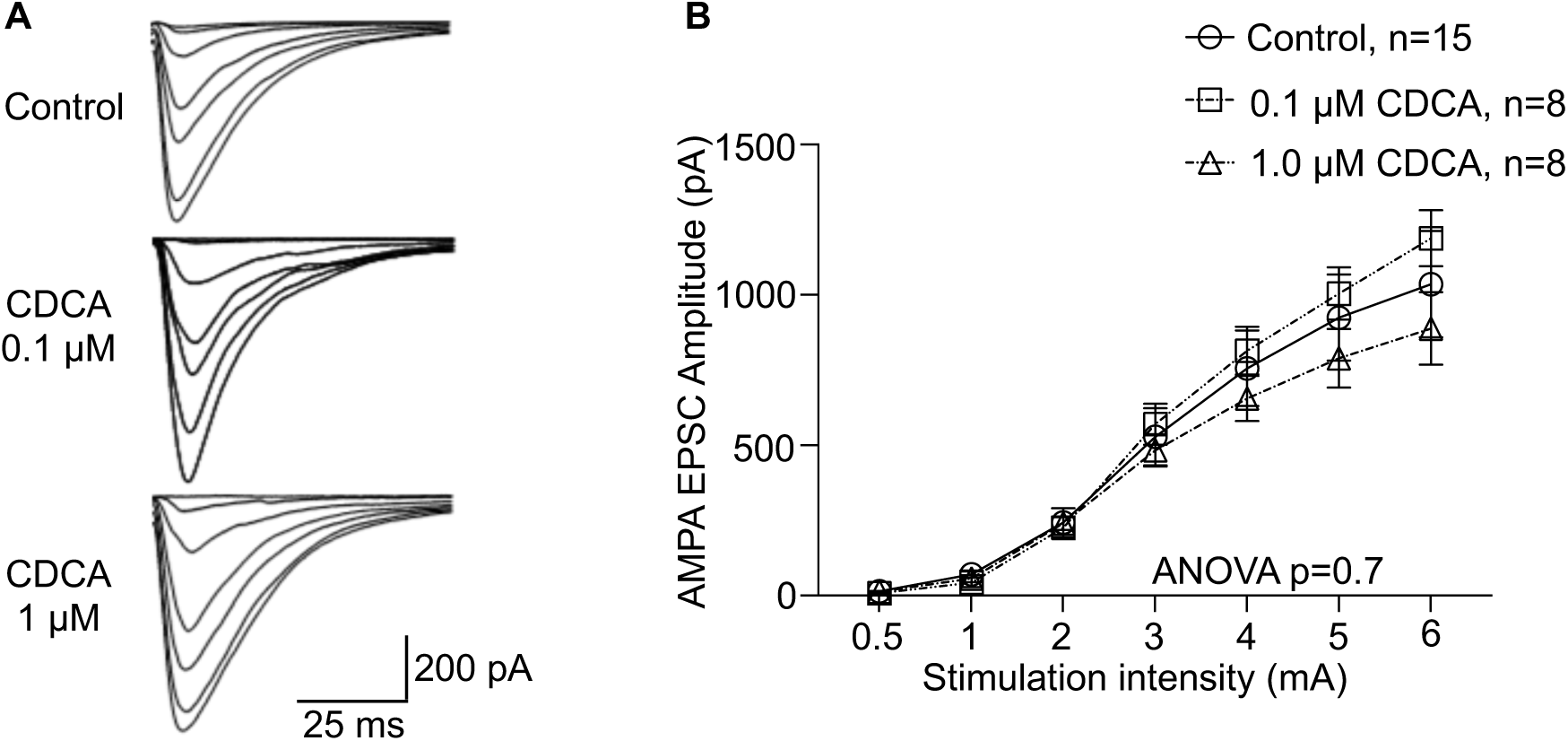
CDCA does not significantly modulate AMPA-mediated excitatory post-synaptic currents. (A) Representative traces of input/output response of AMPAR mediated current recorded from CA1 neurons of acute hippocampal slices from *wild-type* mice treated with CDCA. (B) Quantified input/output responses of AMPAR mediated current. Control, n=15; CDCA 0.1 μM, n=8; CDCA 1 µM, n=8. Data are shown as average ± standard error and are analyzed using Two-way ANOVA followed by Tukey’s multiple comparisons tests.

